# Evolutionary analysis of Trehalose breakdown pathways

**DOI:** 10.1101/2025.10.29.685264

**Authors:** G Ganesh Muthu, Sunil Laxman, Aswin Sai Narain Seshasayee

## Abstract

Trehalose is a widely prevalent, abundant disaccharide that acts as a cellular stress protectant, and functions as an energy source that enters central carbon metabolism when broken down. The evolution and distribution of trehalose breakdown pathways across kingdoms of life have not been studied, and therefore the ability of different organisms to consume trehalose as a carbon source is unknown. In this study, we build a comprehensive evolutionary analysis of the four known trehalose breakdown pathways - trehalase (acid, neutral, glycosyl hydrolase 15), trehalose phosphorylases (TP, treP), and trehalose specific phosphotransferases (PTS), by studying their distributions across ~4000 prokaryotic and eukaryotic genomes. Our study suggests the presence of trehalase in the Last Eukaryotic Common Ancestor (LECA), and reveals near-universal presence of trehalase in eukaryotes, except in all birds where trehalase was lost in the first bird ancestor. Fungi alone retain additional trehalose phosphorylases (TP) in addition to trehalase. In contrast, trehalose breakdown in prokaryotes is highly sporadic but can occur via multiple, independently evolved pathways, including trehalase, the trehalose-specific PTS and trehalose phosphorylase. Finally, we observe that a subset of fast-growing Gammaproteobacteria retain the trehalose specific PTS, the loss of which reduces growth in *Escherichia coli*. Overall, our findings uncover the evolutionary landscape of trehalose breakdown, and use of this versatile disaccharide as an energy reserve in different kingdoms of life.

## Introduction

Sugars are the cornerstone of cellular bioenergetics, and the evolution of metabolic pathways for sugar utilization have enabled organisms to exploit diverse energy sources. Glycolysis is a nearly universal, ancient pathway for glucose metabolism, and multiple sugar breakdown pathways feed glycolysis. Interestingly, the strategies for breaking down more complex sugars, like the non-reducing disaccharide trehalose, are remarkably diverse. Trehalose is an abundant, widely-prevalent disaccharide with unique biochemical properties due to its stable α,α-1,1 glycosidic bond that holds two glucose molecules together. This stability allows it to function as a versatile protectant against desiccation, heat, cold and osmotic stress, and as a signaling molecule (Paul et al., 2008, Iturriaga et al., 2009, Erkut et al., 2011). This makes it a preferred storage carbohydrate, a “universal protectant” and a “chemical chaperone,” for organisms facing periods of metabolic dormancy or environmental adversity (Crowe., 2007). Interestingly, trehalose can also rapidly be enzymatically hydrolyzed into glucose, providing a ready reserve of energy that fuels glycolysis, balances carbon metabolic flux and acts as an energy reserve (Shi et al., 2010, Varahan et al., 2019, Varahan & Laxman., 2021, van Heerden et al., 2014, Yan et al., 2021, Gupta and Laxman 2021, Matsushita & Nishimura, 2020).

While trehalose utilization has been reported across various life forms, its breakdown and metabolism exhibit considerable diversity. Four distinct enzymatic pathways have been identified to break down and utilize trehalose as an energy source (Fig 1A) (Sakaguchi et al., 2020, Shrestha et al., 2024, Boos et al., 1990). The direct hydrolysis of trehalose by trehalase (Treh) results in two glucose molecules, which can directly therefore enter glycolysis. However, alternative pathways to breakdown exist. Trehalose phosphorylase (TP), catalyzes the reversible phosphorolysis of trehalose into glucose-1-phosphate and glucose. This reversibility allows the enzyme to function in both the synthesis and catabolism of trehalose, providing metabolic flexibility (Schwarz A et al., 2007, Shrestha et al., 2024). Another pathway utilizes a phosphoenolpyruvate-dependent phosphotransferase system (PTS) to import and phosphorylate trehalose, which is then hydrolyzed by trehalose-6-phosphate hydrolase (TPH) to produce glucose and glucose-6-phosphate (Escalante et al., 2012). These pathways to breakdown trehalose, and their distinct products, are illustrated in Fig. 1A.

**Figure 1:**
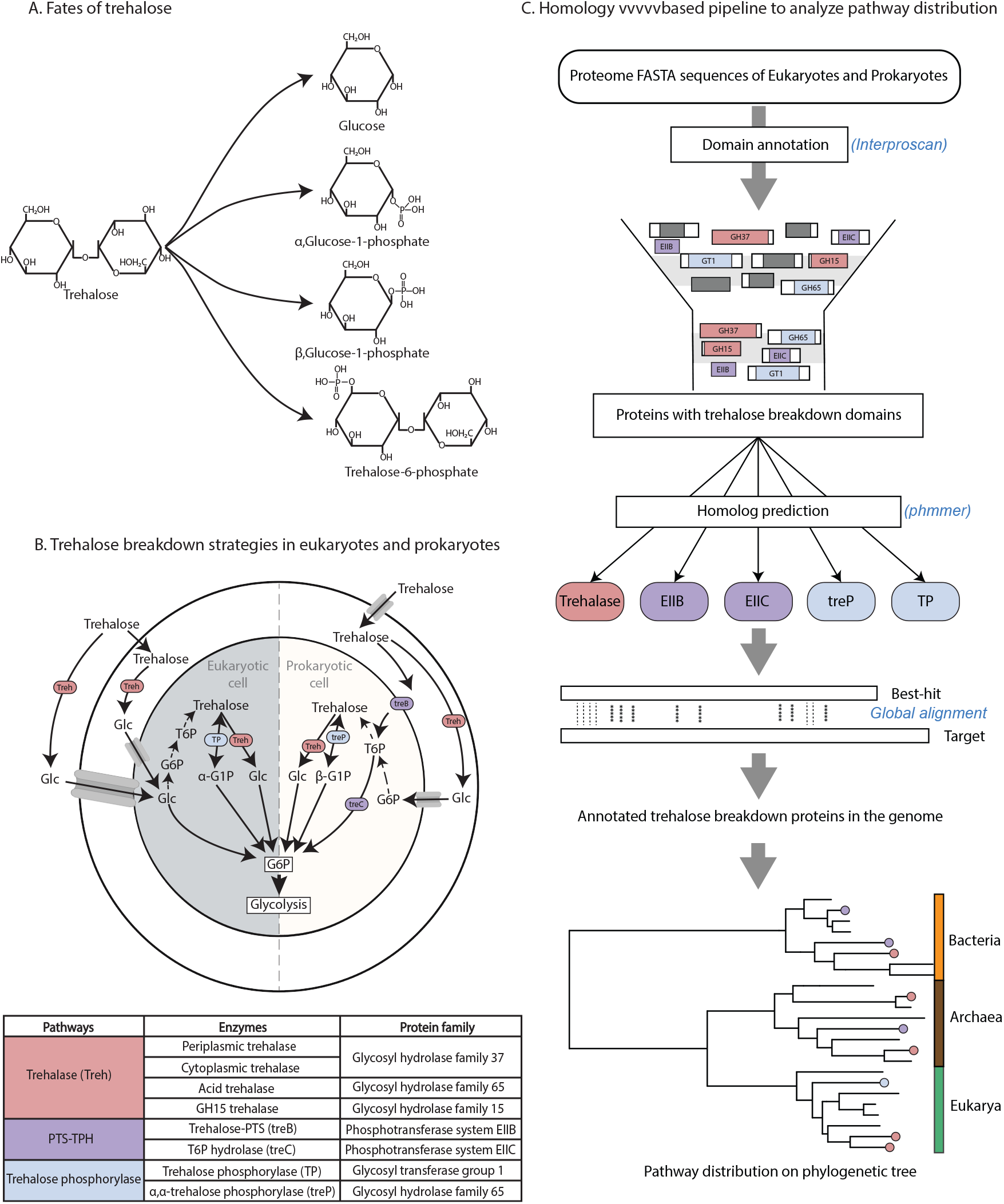
Trehalose breakdown to multiple metabolic intermediates via different pathways: (A) Schematic representation of the four known trehalose breakdown pathways identified in different organisms. Trehalose can be hydrolyzed or phosphorylated to yield glucose or glucose-6-phosphate, which subsequently enter glycolysis for energy generation.(B) Illustration depicting the trehalose breakdown pathways in eukaryotic (left) and prokaryotic (right) cells. In both systems, trehalose breakdown produces glucose or glucose-6-phosphate that feeds into glycolysis for energy generation. The accompanying table summarizes each pathway, the key enzymes, and their corresponding protein family classifications. (C) Overview of the homology-based bioinformatics pipeline used to identify and map trehalose breakdown pathways across 2,583 bacterial, 62 archaeal, and 868 eukaryotic genomes

However, despite our mechanistic understanding of these individual enzymes, a comprehensive picture of their evolutionary origins, and their distribution across the tree of life is unknown. When grouped by known distributions, we can currently only generally illustrate their prevalence in bacterial or eukaryotic cells (as summarized in Fig. 1B). In this study, we therefore asked how the different strategies for trehalose utilization may have evolved. To address this, we performed a large-scale comparative genomic analysis of over 3,800 prokaryotic and eukaryotic genomes, mapping the distribution of trehalose breakdown and reconstructing their evolutionary histories. Our results reveal a clear dichotomy in trehalose utilization between prokaryotes and eukaryotes. Most eukaryotes have retained a trehalase-based system to break down trehalose to glucose, with a notable exception in birds. In contrast, the distribution of these pathways among prokaryotes is sporadic, shaped by horizontal gene transfer, with unique associations specifically in fast-growing Gammaproteobacteria. This work establishes a comprehensive evolutionary framework for understanding how diverse organisms have adapted distinct strategies to break down and utilize trehalose as an energy reserve, and can aid in the informed metabolic engineering of industrially important strains and microbes.

## Results and Discussion

### Dataset description and pathway prediction

We analyzed the distribution of trehalose breakdown pathways across 2,583 bacterial, 62 archaeal, and 868 eukaryotic genomes using a homology-based approach.

We searched for four major trehalose catabolic pathways, comprising six key enzymes, distributed across all domains of life (Fig. 1A, B). We used a total of six KEGG Ortholog (KO) groups corresponding to these enzymes as references for sequence searches. Three types of trehalase enzymes have been identified across different taxa — neutral trehalase (Treh^N^), acid trehalase (Treh^A^), and glycosyl hydrolase family 15 trehalase (GH15-Treh) (Sakaguchi et al., 2020). Treh^N^, a cytosolic and periplasmic enzyme belonging to the GH37 family, functions optimally at neutral pH and hydrolyzes intracellular trehalose. In contrast, Treh^A^ is a GH65 family enzyme localized to vacuoles or the cell surface, where it acts at acidic pH to hydrolyze extracellular trehalose. Despite these distinctions, both Nth and Ath fall under the same KEGG ortholog (K01194), whereas GH15-type trehalase is categorized separately (K22934) (Sakaguchi et al., 2020). Two types of trehalose phosphorylase enzymes have been described in the literature: one specific for the α-anomeric form and another for the β-anomeric form of glucose and glucose-1-phosphate (Shrestha et al., 2024). These enzymes catalyze distinct reactions and correspond to two separate KEGG orthologs: K22248 for the α-specific (retaining) TP and K05342 for the β-specific (inverting) treP, the latter reported exclusively in certain bacteria (Van der Borght et al., 2011). The trehalose-specific phosphotransferase system (PTS) responsible for trehalose import and phosphorylation is represented by K02819, while the downstream trehalose-6-phosphate hydrolase, which converts trehalose-6-phosphate to glucose and glucose-6-phosphate, corresponds to K01226. An organism was considered to possess a trehalose breakdown pathway if its genome encoded all the essential enzymes for that specific pathway.

In summary, we identified 3378 proteins involved in trehalose breakdown across eukaryotes and prokaryotes. We used this dataset for all the analyses described below.

### Eukaryotes except birds retain trehalose breakdown systems

We analyzed the distribution of all trehalose breakdown pathways across 868 eukaryotic genomes and mapped them onto a phylogenetic species tree. Overall, 76% of eukaryotic organisms (660/868) possess at least one trehalose breakdown pathway (Fig. 2A, B). Trehalase (Treh) is the predominant trehalose breakdown enzyme, present in 85.1% (562/660) of trehalose-utilizing eukaryotes (Fig. 2A, B). We carried out a maximum-likelihood-based ancestral state reconstruction of trehalase presence on the eukaryote tree and found that the Last Eukaryotic Common Ancestor likely encoded a neutral trehalase (Treh^N^) (Fig. 2C). Thus, Treh^N^ represents the primary and ancestral mechanism of trehalose catabolism in eukaryotes. In contrast, acid trehalase (Treh^A^), which functions in extracellular trehalose utilization, is restricted largely to fungi (128/193 species), with the exception of a few Basidiomycetes and Sordariomycetes that lack it (Supplementary Fig. 1A).

**Figure 2:**
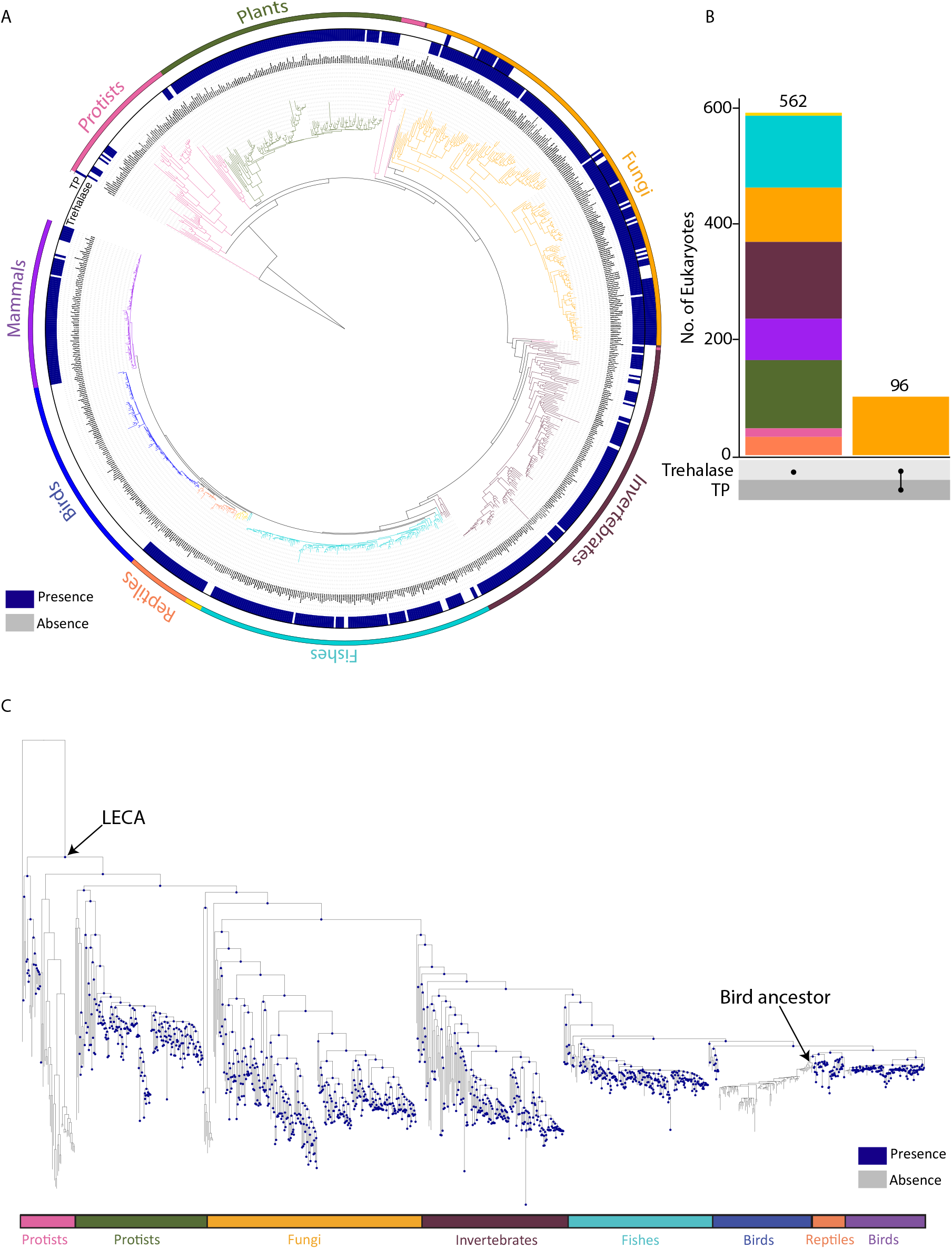
Trehalose breakdown by trehalase is ubiquitous in eukaryotes with a complete loss in birds: (A) Eukaryotic species phylogeny with a heatmap showing presence (blue) and absence (white) of trehalase (Treh) and trehalose phosphorylase (TP) pathways across 868 genomes. (B) UpSet plot showing the combinations of pathway presence across eukaryotic clades, highlighting the dominance of trehalase as the primary trehalose catabolic route. (C) Maximum-likelihood ancestral state reconstruction of trehalase presence across the eukaryotic phylogeny. The Last Eukaryotic Common Ancestor (LECA) most likely encoded Treh, which was vertically inherited in most lineages but lost in the avian ancestor. Tip and node colors indicate trehalase presence (blue) when posterior probability >70%.

Notably, trehalase genes were completely absent across the entire clade of birds (Fig. 2A, C). The avian common ancestor lacked Treh^N^, and it has not been regained in any subsequent bird lineage (Fig. 2C). This is consistent with Brun et al. (2022), who found no evidence of the gene for Treh in a smaller set of bird genomes, and no detectable enzymatic trehalase activity in multiple bird tissue extracts. Their synteny analyses further suggest that Treh^N^ was lost via a genomic inversion event after the crocodile–bird split.

The loss of Treh is particularly striking given that many birds consume trehalose-rich diets (e.g., insects, seeds). This suggests birds may rely on alternative strategies for trehalose metabolism: possibly uptake mediated through symbiotic gut microbiota, or other non-*Treh*-based mechanisms. Brun et al. have speculated that low levels of trehalose breakdown activity they observe in birds might arise from the microbiome. We therefore analyzed gut microbiota from 15 bird species spanning multiple clades. These microbial communities encoded several trehalose breakdown pathways, including bacterial PTS-TPH, trehalase, and treP (Supplementary Fig. 1B). However, the bird gut microbiome is not particularly unique in this respect; analysis of gut microbiomes of mammals and reptiles show similar occurrences of these pathways (Supplementary Fig. 1C). Interestingly, 36 of the 91 bird genomes analyzed also encode a TrehA-like protein belonging to the GH65 family, resembling the fungal acid trehalase (Supplementary Fig. 1A). However, these homologs lack the N-terminal domain required for cell-surface localization, and their functional relevance in non-fungal eukaryotes remains uncertain. Together, these results indicate that the evolutionary loss of Treh in birds is likely irreversible in birds, and that their gut microbiota might only partially compensate for dietary trehalose intake.

Beyond birds, trehalase is also sporadically absent in a few mammals including the gray seal (*Halichoerus grypus*), dromedary camel (*Camelus dromedarius*), and several cetaceans such as dolphins and whales (Fig. 2A). This observation is consistent with the genome-wide study showing that the trehalase gene (Treh) has been independently lost or pseudogenized across some mammalian lineages, and which may be linked to changes in dietary composition (Jiao et al., 2019).

In addition to trehalase, we identified the trehalose phosphorylase (TP) pathway, which catalyzes the reversible conversion of trehalose into α-glucose-1-phosphate and glucose (Fig 1B), as the only other trehalose breakdown pathway present predominantly in fungi. TP is found across several fungal lineages, including Basidiomycetes such as *Agaricomycetes, Tremellomycetes*, and *Rhodotorula*, and is also widespread in Ascomycetes (Fig. 2A, B). This substantially expands upon earlier reports describing TP activity in fungi such as *Agaricus bisporus* (Wim et al., 1998) and *Pichia fermentans* (Schick et al., 1995). Note here that the TP enzyme, unlike Treh, can also be used for trehalose synthesis; it is however not known whether its primary function is restricted to breakdown, or for synthesis as well. Nearly all fungi harboring the TP pathway also encode the trehalase enzyme (which exclusively breaks down trehalose), with one exception—*Arthroderma uncinatum* (Saccharomycetes)—which contains TP but lacks trehalase. Notably, Ascomycetes represent the most metabolically versatile group, frequently encoding neutral trehalase (Treh^N^), acid trehalase (Treh^A^), and trehalose phosphorylase (TP) within the same genome (Fig. 2A, B; Supplementary Fig. 1A). This combination highlights the multifaceted role of trehalose breakdown in these organisms, as a readily accessible energy source that can re-enter carbon metabolism.

Finally, trehalose breakdown pathways are relatively uncommon among protists, with only 15 of the 71 species analyzed possessing the trehalase pathway. Despite this rarity, trehalose utilization spans a wide ecological range, occurring in both parasitic and free-living protists. These include organisms from the SAR (Stramenopiles, Alveolates, Rhizaria) lineage, such as Phytophthora, Saprolegnia (both parasitic) and Paramecium (free-living), as well as Dictyostelium, which belongs to the Amoebazoa. Notably, excavate parasites such as Trypanosoma and Leishmania completely lack trehalose breakdown pathways (Fig. 2A), suggesting that these organisms do not utilise trehalose and only depend on alternative carbon sources and metabolic strategies.

Together these findings show that trehalase is a dominant and ancestral trehalose breakdown enzyme in eukaryotes, retained in most lineages but lost entirely in birds and sporadically in select mammalian clades. Protists exhibit only sporadic presence of trehalose metabolism. Fungi alone (amongst eukaryotes) display additional diversity with the presence of the TP pathway alongside (often multiple) trehalase enzymes. Collectively, these data suggest the fairly ubiquitous use in eukaryotes of trehalose as an energy reserve (via trehalase activity), with clear exceptions.

### Trehalose breakdown pathways are sporadically distributed in prokaryotes

While analysing the distribution of trehalose breakdown pathways in the gut microbiomes of birds and other animals, we noticed multiple bacterial mechanisms for trehalose breakdown. This prompted us to examine how widespread these pathways are across all prokaryotes, and we therefore assessed the presence of all known trehalose breakdown pathways in prokaryotes. Extending our analysis to 2,645 bacterial genomes, we found that only 35% of species (931/2,645) encoded one or more trehalose breakdown pathways (Fig. 3A, B), suggesting that the ability to break down trehalose is relatively uncommon among prokaryotes.

**Figure 3:**
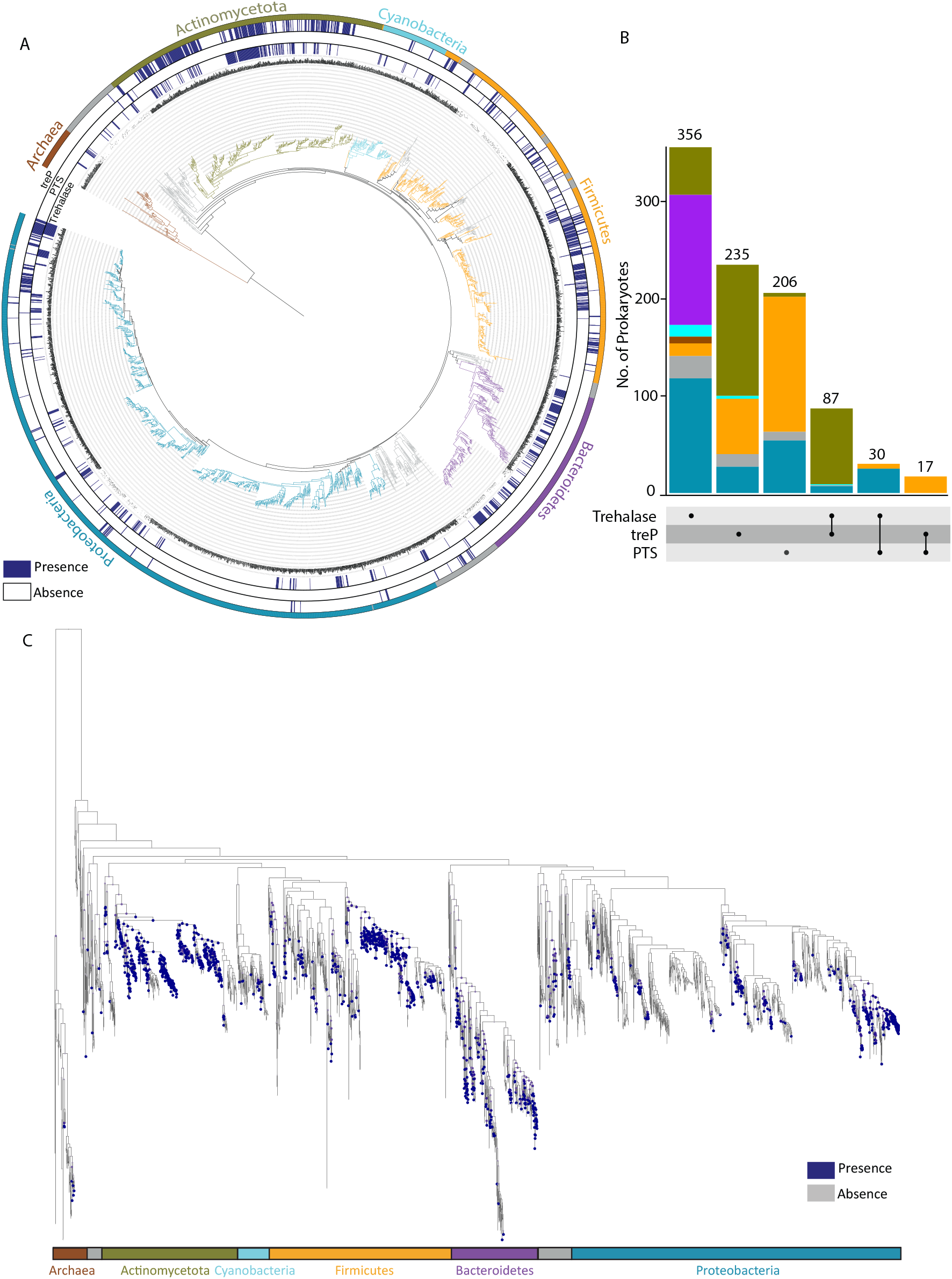
Trehalose breakdown pathways are sporadically distributed and acquired independently across prokaryotes: (A)Phylogenetic tree of 2,645 prokaryotic species showing the distribution (presence in blue, absence in white) of trehalase (Treh), trehalose-specific phosphotransferase system (PTS), and trehalose phosphorylase (treP) pathways. (B) UpSet plot summarizing combinations of trehalose breakdown pathways across bacterial phyla, illustrating the mosaic distribution. (C) Maximum-likelihood-based ancestral state reconstruction of overall trehalose breakdown pathways in prokaryotes. Trehalose breakdown pathways are gained and lost multiple times through evolution. Node colors represent the probability of pathway presence (blue; posterior probability >70%).

Among the four pathways identified, Treh is the most widespread, present in 50.8% (473/931) of trehalose-utilizing organisms (Fig. 3A, B). Of these, the neutral trehalase (Treh^N^) is the predominant Treh enzyme for trehalose catabolism in these taxa (67.4%; 319/473), with a strong representation in Bacteroidetes (134/294), Actinomycetota (127/445), and Proteobacteria (148/1130). The acid trehalase (Treh^A^), in contrast, was sporadically distributed, occurring only in 154 bacterial species, with notable presence in certain Bacteroidetes (52/294) where it often co-occurs with Treh^N^ (Supplementary Fig. 2A). Within Bacteroidetes, trehalose catabolism appears to rely exclusively on trehalase activity— either Treh^N^, Treh^A^, or both (Supplementary Fig. 2A).

Interestingly, archaea are largely devoid of trehalose breakdown pathways. In our dataset, only 7 of 69 species had any trehalose breakdown pathway (exclusively Treh), and a similar pattern was observed in a larger dataset of 264 archaeal genomes, where only 25 species harbored trehalose breakdown capacity (Supplementary Fig. 2B). The few archaea capable of trehalose catabolism all encode a Treh enzyme of the Glycosyl Hydrolase (GH) 15 family. This class of Treh has been previously described in thermophilic archaea (Lee et al., 2018; Sakaguchi et al., 2015) as well as in extremophilic bacteria (Zhang et al., 2022), where it functions in trehalose hydrolysis under extreme environmental conditions. Consistent with this, we find GH15 trehalases mainly in archaea belonging to the Methanomicrobia and Halobacteriales, two groups well adapted to extreme habitats such as high salinity and anaerobic environments (Supplementary Fig. 2B inset).

Under low osmolarity conditions, trehalose can be imported into the cell through the trehalose-specific *treB*-encoded phosphotransferase system (PTS). During transport, trehalose is phosphorylated to trehalose-6-phosphate and subsequently hydrolyzed by the *treC*-encoded enzyme to yield glucose and glucose-6-phosphate (Boos et al., 1990). This pathway, previously reported in a few bacteria for intracellular trehalose utilization (Wu at al., 2023, Park et al., 2016), was detected in 253 bacterial species, with notable representation in Firmicutes (160/577) and Gammaproteobacteria (75/337) (Fig. 3B, C). Although not widespread, its presence across phylogenetically diverse taxa suggests that the PTS-TPH system might enable these bacteria to import and utilize extracellular trehalose as an additional carbon and energy source.

We further identified the bacterial specific trehalose phosphorylase (treP) pathway in 359 species, particularly in Actinomycetota (213/445; 47.8%), Firmicutes (74/577) and Cyanobacteria (4/116). Note here that the bacterial treP belongs to a different enzyme family from its fungal relative (TP).

We carried out an ancestral state reconstruction of trehalose breakdown pathways in bacteria and archaea. This revealed a sporadic pattern of gains of the respective pathway (Fig. 3C). Independent acquisitions were inferred at the ancestral nodes of Actinomycetota and Firmicutes, while most other gains occurred at terminal, lineage-specific branches of the phylogeny. Independent gains were also identified in the ancestor of the Halanaerobiales order, and in a later Firmicutes ancestor that gave rise to the Lactobacillales and Bacillaceae families. We observed scattered gains of trehalose breakdown pathways in groups such as Cyanobacteria, Spirochaetes, Bacteroidetes, and Proteobacteria. These findings collectively establish that the emergence of trehalose utilization in prokaryotes was not the result of a single ancestral innovation, but likely occurred multiple times independently across diverse clades.

In summary, in contrast with eukaryotes, trehalose breakdown is present in only about one-third of prokaryotes, with clear phylum-specific differences. However, bacteria exhibit a large diversity of distinct trehalose breakdown pathways. Notably, Actinomycetota are particularly well equipped (two-thirds of genomes encoding these pathways), often encoding multiple trehalose breakdown pathways, Bacteroidetes rely solely on Treh, and Firmicutes and Proteobacteria retain both the PTS and treP systems. Trehalose metabolism in prokaryotes is not an ancestral, conserved trait but rather specific to particular lineages.

### Extensive Horizontal Gene Transfer of Trehalose Breakdown Pathways in Prokaryotes

To test the extent to which the distribution of trehalose breakdown pathways in prokaryotes reflects phylogenetic relationships, we calculated the phylogenetic signal for each pathway using the delta statistic (Borges et al., 2019). This method compares the observed distribution of traits across a phylogeny, to a null expectation generated by randomizing trait presence/absence while keeping the total number of presences constant. The delta statistic measures Shannon entropy of ancestral state reconstructions, where lower entropy indicates stronger phylogenetic signal (i.e., the trait tracks the phylogeny more closely). Our analysis revealed only a weak phylogenetic signal for all trehalose breakdown pathways, with mean delta values of 5.37 (trehalase), 10.45 (PTS), and 7.88 (treP). In contrast, the control gene (cytochrome oxidase COX1), present in 64.25% of prokaryotes in our dataset, exhibited a strong phylogenetic signal (mean delta = 36.43) (Supplementary Fig. 3).

To investigate the influence of horizontal gene transfer (HGT) in the observed distribution of pathways, we employed phylogenetic reconciliation of gene trees within a species tree using a parsimony-based Duplication-Transfer-Loss (DTL) framework. In this method, evolutionary events are assigned specific costs, and the reconciliation aims to minimise the total cost. This analysis revealed that ~43% of all extant trehalose breakdown pathways are downstream of a horizontal gene acquisition event (Fig. 4).

**Figure 4:**
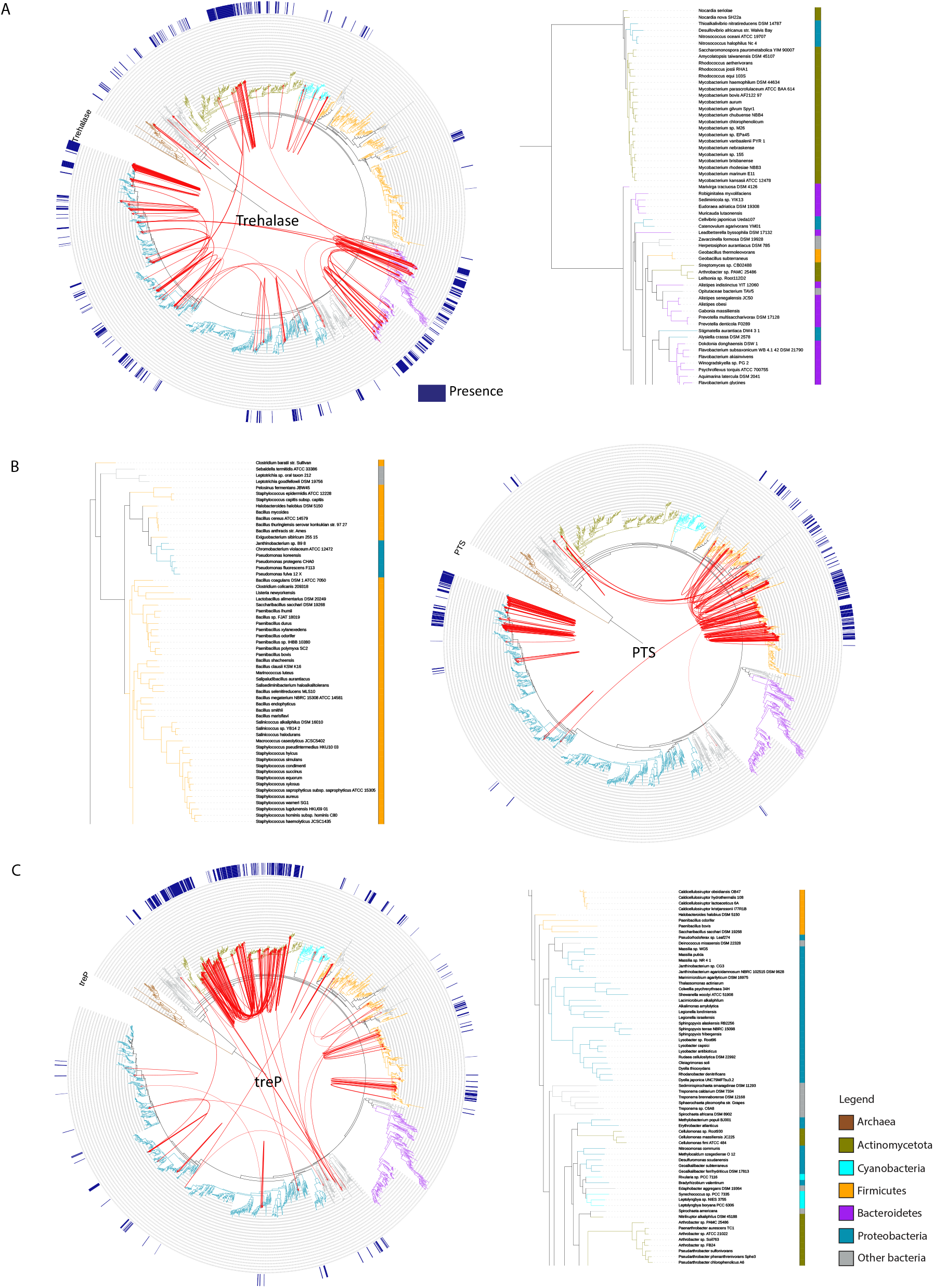
Extensive horizontal gene transfer (HGT) shapes the landscape of trehalose breakdown pathways in prokaryotes: Species trees showing inferred HGT events for (A) Treh (trehalase), (B) PTS, and (C) treP genes based on gene–species tree reconciliation using the Duplication–Transfer–Loss (DTL) model. Arrows denote frequent donor–recipient clades for each gene. Inset panels display corresponding gene trees, showing cross-phyla clustering consistent with HGT events (See Supplementary material for exact number of reconciliations for each donor-recipient pair).

At the pathway level, we found distinct patterns of transfer for each enzyme/pathway:

For the Treh genes, Proteobacteria emerged as both donors and recipients, with multiple intra-clade transfers as well as inter-clade transfers involving Actinomycetota and Bacteroidetes. Gene-tree of trehalase show clustering of certain Proteobacterial trehalases with Actinomycetotal and Bacteroidetes homologs, consistent with cross-phyla exchange (Fig 4A inset).

For the PTS pathway, gene transfers were largely restricted to within Firmicutes. Only rare cross-phyla events were observed, including *Clostridium innocuum* donating to a Gammaproteobacterium (*Cardiobacterium valvarum*) and *Clostridium lentocellum* donating to Actinomycetota (*Coriobacterium glomerans*) (Fig. 4B inset).

For the TP genes, Actinomycetota appear to be the major donors, with gene transfers to Firmicutes, and Proteobacteria. This broad dissemination suggests that treP might be a particularly mobile pathway that could provide a species unique advantage. Finally, for the GH15 archaeal trehalases, these genes are found exclusively in archaea, and appear to have originated from Actinomycetota, with subsequent transfers occurring within archaeal clades (Supplementary Fig. 4A).

Collectively, these extensive analyses suggest the independent emergence of, and evidence of extensive horizontal gene transfer in shaping the landscape of trehalose breakdown in prokaryotes.

### The trehalose specific PTS pathway is found in fast-growing Gammaproteobacteria

Trehalose breakdown through the various pathways discussed in this study yields glucose or glucose-6-phosphate, which feeds directly into glycolysis-the primary route for energy generation and biomass production. The presence of more than one trehalose breakdown pathway within a genome could therefore aid metabolic flexibility, allowing organisms to efficiently mobilize trehalose under different environmental conditions. Redundancy in trehalose catabolism may facilitate the supply of glycolytic intermediates, potentially supporting higher growth in specific environments. Therefore, we asked whether the ability to breakdown and utilise trehalose had any associations with bacterial growth. Across bacteria, the number of 16S rRNA operons serves as a robust genomic proxy for growth rate, as species with higher copy numbers can initiate protein synthesis more rapidly and respond swiftly to nutrient availability (Vieira-Silva & Rocha, 2010; Couturier & Rocha, 2006). Given that trehalose functions as an energy reserve, we asked if the ability to break down trehalose in bacteria had any associations with growth rate. To test this, we examined the relationship between trehalose breakdown pathways and 16S rRNA gene copy numbers across bacterial genomes.

First, we assessed if all the trehalose-utilizing bacteria encode the key glycolytic enzymes pyruvate kinase and GAPDH, in order to confirm their ability to funnel trehalose-derived sugars to fuel glycolysis. We found that genomes encoding Treh or the PTS-TPH pathway universally contained GAPDH, and all but one also contained pyruvate kinase (Supplementary Fig. 5A,B).

We then assessed 16S rRNA gene copy numbers between bacteria that are trehalose utilizers and those that are non-utilizers. Overall, we observe that trehalose-utilizing organisms displayed significantly higher 16S rRNA copy numbers (Fig. 5A). Among the pathways, the PTS-TPH pathway was significantly associated with bacteria that have higher growth rates (Fig. 5A, inset). However, when analysed on a dataset comprising all bacteria without any phylogenetic stratification, trehalose-PTS showed no significant difference in growth rate when compared to bacteria encoding other sugar PTS systems (Supplementary Fig. 4B).

**Figure 5:**
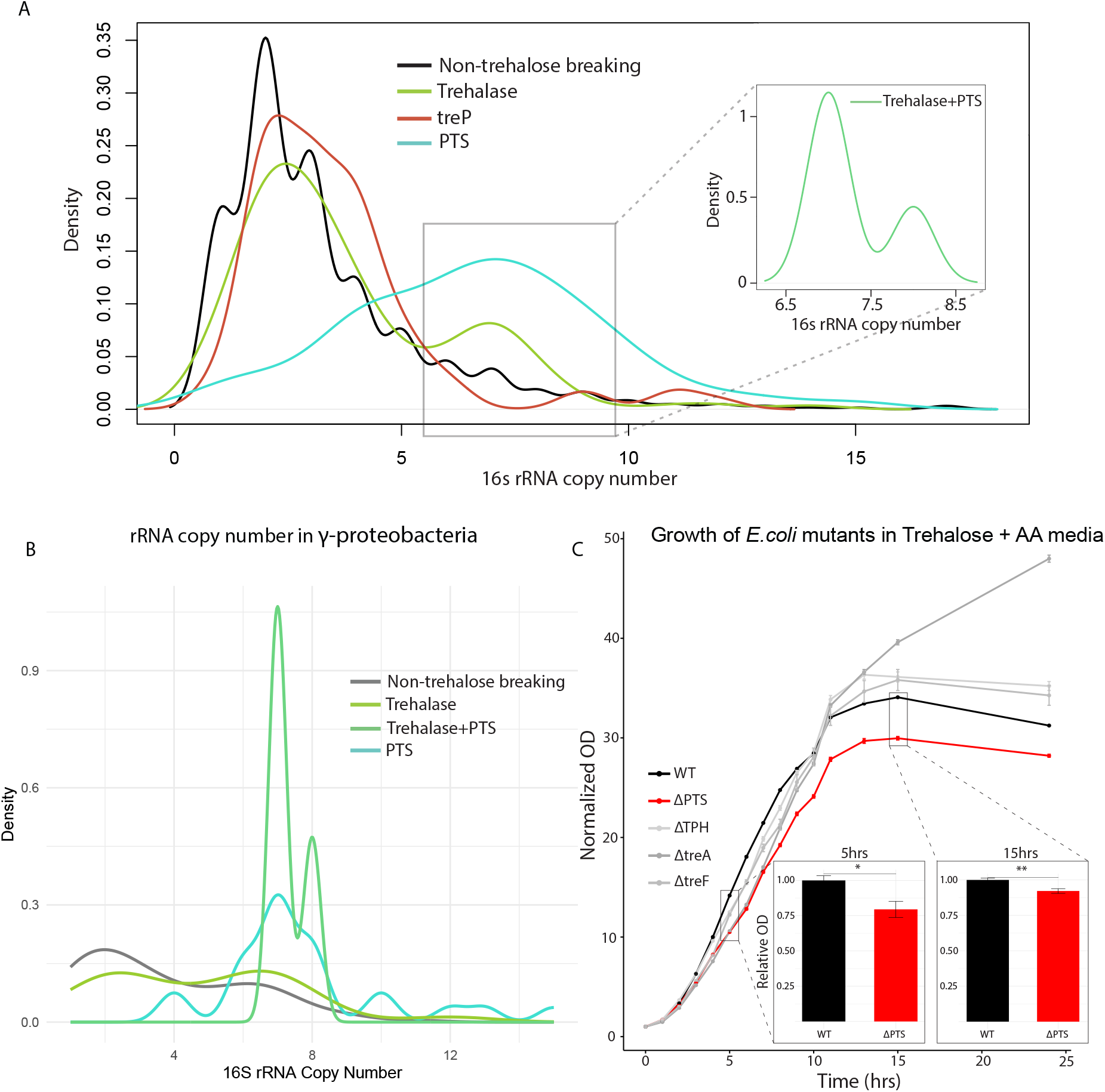
Trehalose-specific PTS is associated with fast growing γ-proteobacteria: (A) Density plot of 16S rRNA gene copy numbers—used as a proxy for maximal growth rate—comparing trehalose-utilizing and non-utilizing bacteria. The inset highlights that genomes encoding both trehalase and the PTS-TPH pathway show significantly higher predicted growth rates (Wilcoxon rank-sum test: non-utilizers vs PTS, p = 6.33×10^−19^; non-utilizers vs trehalase, p = 6.91×10^−04^; non-utilizers vs treP, p = 0.0144). (B) Within γ-Proteobacteria, strains encoding PTS-TPH and those encoding both PTS-TPH and trehalase show significantly elevated 16S rRNA copy numbers relative to non-utilizers (Wilcoxon rank-sum test: non-utilizers vs PTS+trehalase, p = 2.96×10^−08^; non-utilizers vs PTS, p = 8.09×10^−14^). (C) Growth curves of E. coli wild-type (WT) and knockout strains lacking treB (PTS), treC (trehalose-6-phosphate hydrolase), treA (periplasmic trehalase), or treF (cytoplasmic trehalase) cultured in M9 minimal medium supplemented with amino acids and trehalose as the sole carbon source. Data represent mean ± SD from three biological replicates. Insets show relative OD_600_ of WT versus ΔtreB at 5 h and 15 h (two-tailed t-test: 5 h, p = 0.0103; 15 h, p = 0.003667). Asterisks indicate statistically significant differences.

We therefore investigated what this subset of fast-growing bacteria, which encoded both Treh and PTS-TPH, were. We noted that these genomes belonged exclusively to the Gammaproteobacteria (Fig. 5A, inset). We therefore further investigated this clade of bacteria more closely, by comparing with Gammaproteobacteria that do not contain the Treh and PTS-TPH genes. We observe that within Gammaproteobacteria, genomes that encode PTS-TPH have higher 16s rRNA copy numbers/faster growth rates than those (a) without any trehalose breakdown pathways, or (b) with alternate PTS enzymes that are specific for sugars other than trehalose, or (c) with Treh alone (Fig. 5B). The other clades of bacteria with multiple trehalose breakdown pathways are the Firmicutes and Actinomycetota. Notably, such a trend and association of the trehalose PTS system was not observed in these bacteria (Supplementary Fig. 5C,D). Collectively, these analyses reveal that the presence of trehalose PTS may be associated with faster growth in Gammaproteobacteria (although we cannot rule out confounding phylogeny, as Gammaproteobacteria with PTS-TPH are often Enterobacteriaceae).

These results suggest the possibility of the utilization of the PTS system to support faster growth in Gammaproteobacteria. In order to test whether PTS-TPH might support higher growth we used *E. coli* (a model Gammaproteobacterium), and assayed growth of knockout strains lacking *treB* (PTS), *treC* (trehalose-6-phosphate hydrolase), *treA* (periplasmic trehalase), or *treF* (cytoplasmic trehalase), using media containing only amino acids and trehalose as carbon source. All the trehalose breakdown enzyme knock-outs show a decreased initial growth rate (apparent after 4 hours of growth). Notably, a clear growth reduction is observed continuously over ~24 hours of growth in the Δ*treB* (PTS deficient) *E. coli* strain compared to the wild type (WT) strain (Fig. 5C, Supplementary Fig E, F, G). Interestingly, the Δ*treA* (periplasmic trehalase) mutants grew faster than WT during the later growth phase (~13 hours; Fig 5C, Supplementary Fig. 5G). This result may be explained by experimentally established regulatory effects, where the loss of *treA* leads to the induction of the sugar transporters like *treB*, and enhances trehalose uptake and utilization (by the PTS system) (Shimada et al., 2011; Jakowec et al., 2023). These data therefore reveal that the fast growing Gammaproteobacteria might utilize the PTS trehalose breakdown system to their advantage, to support their high growth rates in trehalose-replete environments.

## Conclusion

This work provides a comprehensive evolutionary overview of trehalose breakdown across all domains of life. We show that eukaryotes are strongly committed to trehalose catabolism and the use of trehalose as an energy reserve, with trehalase as the dominant and ancestral enzyme conserved across most lineages—except in birds. Its broad conservation across eukaryotes, with an overrepresentation in fungi, underscores the metabolic importance of trehalose as a mobilizable carbon and energy source. The biochemical importance of trehalase, which is abundant in mammalian and fungal tissues and insect hemolymph (Shukla E at al., 2015, Tellis et al., 2023), further reflects its physiological significance. In contrast, trehalose breakdown in prokaryotes is sparse and mosaic, shaped by independent evolutionary gains and horizontal gene transfer. Only about a third of bacteria encode these pathways, and fast growing γ-Proteobacteria retain multiple pathways that may support growth in trehalose-rich environments. Archaea largely lack any trehalose breakdown pathways, and therefore likely use trehalose exclusively for protection rather than energy metabolism. This study therefore delineates the evolution of trehalose breakdown with implications towards our understanding of trehalose as a valuable energy reserve to fuel metabolism. The biosynthesis of trehalose —driven by diverse precursors and enzymatic mechanisms—remains more complex, and understanding how these pathways co-evolved with breakdown systems will be crucial to fully define the physiological and evolutionary roles of this versatile disaccharide that serves as a protectant (when present) and an energy source (when broken down).

## Materials and Methods

### Data

Genome assemblies and annotations were retrieved from the NCBI FTP repository (ftp.ncbi.nlm.nih.gov/) using custom automated scripts. Coding nucleotide sequences (.fna) and predicted proteomes (.faa) were downloaded for ~900 eukaryotic and ~12,000 prokaryotic accessions. To generate a non-redundant dataset, one representative genome per species was selected based on the assembly with the highest gene count. The final dataset comprised 868 eukaryotes and 2,645 prokaryotes (2,583 bacteria and 62 archaea), which were used in all downstream analyses. Because archaeal genomes are underrepresented in curated genome sets, an expanded collection of 264 archaeal genomes was additionally downloaded and analyzed separately to validate pathway distribution trends. A unified protein and nucleotide database was constructed by merging all retrieved sequences. For constructing prokaryotic species tree, 16S rRNA sequences were obtained from the Genome Taxonomy Database (GTDB; Parks et al., 2022). Reference protein sequences for enzymes involved in trehalose breakdown pathways were obtained from the KEGG Orthology database (genome.jp/kegg/). The microbiome data of birds, mammals and reptiles are obtained from The Animal Microbiome Database (AMDB) (http://leb.snu.ac.kr/amdb).

### Prokaryotic species phylogeny

For the prokaryotic species tree, one 16S rRNA sequence per genome was retrieved from the GTDB database using an in-house Python script. Multiple sequence alignment (MSA) of the 16S sequences was performed using MUSCLE v5 (Edgar, 2022) with default settings. The alignment with maximum column confidence was taken for further analysis. Before generating the species tree, the sequences were trimmed using trimAL (https://github.com/inab/trimal) with -gappyout for reducing the gaps in the MSA. The trimmed alignment was then passed to IQTREE v2 with model GTR+F+I+R10 (LogL=-478662.3293, BIC=998509.322) which was obtained via model testing (-m TEST) (Hoang et al., 2018; Kalyaanamoorthy et al., 2017; Minh et al., 2020). Three independent runs of IQTREE were run with each having an ultrafast bootstrap of 8,000 and a random seed value of 356708, 16225, and 17225 respectively. The tree that had the highest logLik was taken as the most probable species tree. The resulting tree was rooted with Archaea as the outgroup using the root.tree() function from the phytools v2.3-0 (Revell, 2012) package in R.

### Eukaryotic species phylogeny

For the eukaryotic species tree, an orthologous gene-based approach was employed. Twenty-five out of thirty-seven eukaryotic genes of bacterial ancestry (euBac) reported by (He et al., 2014), were selected for tree construction based on their presence in over 90% of the organisms. Each euBac protein was searched against the protein database using BlastP with an E-value cutoff of 0.01, and the top 500 protein sequences were extracted. MSA for these 500 sequences was performed using MUSCLE v5, and profile HMMs were built using the hmmbuild function of HMMER v3.3.2 (Eddy, 2011). The euBac proteins of eukaryotes and one archaea, *Halobaculum salinum*, which was used to root the phylogenetic tree, were extracted using the hmmsearch function of HMMER v3.3.2, with the profile HMMs as queries. The NCBI protein identifiers in each multi-fasta file were edited to correspond to the organism, and organism name redundancies were removed using an in-house Python script. MSA of each euBac multi-fasta protein file was performed using MUSCLE v5, and the conserved regions relevant for phylogenetic inference were trimmed using BMGE (Criscuolo & Gribaldo, 2010).

The best model for each euBac protein was selected using IQTREE ModelFinder (-m TEST) on each trimmed euBac multi-fasta file. The 25 trimmed euBac multi-fasta files were then concatenated using the Perl script catfasta2phyml.pl (https://github.com/nylander/catfasta2phyml). A gene partition file was created by specifying the start and end positions of each gene after concatenation and providing the best model for each gene. A maximum likelihood phylogenetic tree was then constructed using the concatenated fasta sequences and the gene partition file, with *Halobaculum salinum* designated as the outgroup using the -o option in IQTREE. Branch supports were evaluated using 1,000 ultrafast bootstrap approximations (-bb 1000).

The phylogenetic species trees were visualized and pruned on the iTOL server (Letunic & Bork, 2021). An in-house python script was used to create iTOL datasets for annotating the phylogenetic tree.

### Identification of trehalose breakdown pathways

Protein sequences corresponding to the orthologous groups of enzymes involved in trehalose breakdown pathways—namely, α,α-trehalase (K01194), glycosyl hydrolase (GH) 15 trehalase (K22934), phosphotransferase system (PTS) (K02819), trehalose-6-phosphate hydrolase (TPH) (K01226), α,α-trehalose phosphorylase (K05342), and trehalose phosphorylase (K22248)—were downloaded from the KEGG database as KEGG Ortholog (KO) groups. In total, six KO groups were retrieved, and a local database of trehalose breakdown enzymes was created by combining the protein sequences from all six groups. To annotate the proteomes of each organism, InterProScan v5.36 (Jones et al., 2014) was used to identify protein domains based on the PFAM database. The PFAM domain IDs for domains associated with trehalose breakdown were obtained from the KO groups in the KEGG database (see supplementary material for all the PFAM ids related to each KO group). Proteins containing these specific PFAM domains were extracted from the InterProScan output. Subsequently, a sequence search analysis was performed using phmmer, where each protein was queried against the local KO database of trehalose breakdown enzymes. A global alignment of all hits was conducted against their respective target sequences using the Needleman method of EMBOSS v6.6 (Rice et al., 2000), with a 50% identity threshold applied. Proteins meeting this threshold were then assigned to the appropriate KO group (Fig. 1c).

The presence or absence of trehalose breakdown pathways was determined based on the presence of the KO groups identified in the previous steps. For a pathway consisting of two steps, both KO groups needed to be present for the pathway to be considered present; otherwise, the pathway was deemed absent. This pathway information was then mapped onto the species phylogenetic trees using the iTOL server to visualize its distribution.

### Phylogenetic correlation analysis

To determine whether the distribution of trehalose breakdown pathways is correlated with phylogeny, we performed a phylogenetic correlation analysis using the delta statistic (Borges et al., 2019). This analysis was conducted using the source code provided by Borges et al. 2019 and using the phytools v2.3-0 package in R. The delta statistic compares the observed distribution of traits across a phylogeny to the distribution expected under a null model calculated by randomizing the presence and absence of a trait.

The species tree was rooted at the midpoint, and the delta statistic was calculated for each pathway across 100 iterations. The observed data were randomized, maintaining the number of presences and absences constant, and the delta statistic was recalculated as an internal control. The mitochondrial gene COX1 was used as an additional control. The distribution of delta values for both observed and randomized data was then visualized using the ggplot2 v3.5.1 package in R.

### Ancestral state reconstruction analysis

To trace the evolutionary history of trehalose breakdown pathways, ancestral state reconstruction of the KO groups involved in these pathways was performed using the phytools v2.3-0 package. Two distinct character states were defined for each KO group: presence and absence. The species phylogenetic trees were rooted at the midpoint using the midpoint.root() function. Ancestral character states at each node were estimated through stochastic character mapping, with 100 simulations performed using the make.simmap() method. The posterior probability (pp) of each character state at each node was calculated and compared across three discrete character evolution models: Equal Rates (ER), Symmetric (SYM), and All Rates Different (ARD). Based on Akaike Information Criterion (AIC) weights, the ER model was selected as the best-fitting model. The posterior probabilities at each node that are greater than 0.7 are considered as presence and were then plotted onto the midpoint-rooted phylogenetic tree using the ggtree (Yu et al., 2017) package in R.

### Horizontal gene transfer analysis

RANGER-DTL 2.0 (Bansal et al., 2018) was employed to predict the donors and recipients of horizontal gene transfer (HGT) events among and within bacterial and archaeal species. The prokaryotic species tree, constructed using 16S rRNA with archaea as the outgroup, served as the reference species tree for this analysis. The protein sequences predicted using HMMER for each KO group were used to construct individual gene trees for the respective KO group. MSA was done using MUSCLE which was used as the input for IQTREE with the ModelFinder option (-m TEST) to choose the best model and with an ultrafast bootstrap of 1000. Optimal rootings for these gene trees were determined using the OptRoot program with default settings to minimize the Duplication–Transfer–Loss (DTL) reconciliation cost. The RANGER-DTL program was then used to compute the optimal DTL reconciliation for each rooted species tree and rooted gene tree pair. The analysis was conducted across 100 simulations for each transfer cost (T) value of 1, 2, and 3, resulting in a total of 300 simulations. The AggregateRanger program was subsequently used to compute support values for the most frequently identified donor species from all the 300 simulations with alternative event cost assignments.

An in-house python script was written to extract the nodes with most transfer events and to back trace the most frequent recipient for a given most frequent mapping. The results were viewed as connections from donor to recipient in the species phylogenetic tree on the iTOL server using an in-house python script.

### 16s rRNA gene copy number estimation

For all 2,583 bacterial genomes analyzed, coding nucleotide sequences (.fna) and corresponding genome annotation files (.gff) were downloaded from the NCBI FTP server. The BAsic Rapid Ribosomal RNA Predictor (Barrnap) tool (https://github.com/tseemann/barrnap) was used to identify rRNA genes across all genomes. An in-house Python script was developed to parse Barrnap output and quantify the number of 16S rRNA gene copies per genome.

The distribution of 16S rRNA gene copy numbers was compared between bacterial groups encoding different trehalose breakdown genes. All plots were generated using the ggplot2 package in R. Statistical comparisons between distributions were performed using Wilcoxon rank-sum tests.

### Bacterial strains and growth curve assay

Escherichia coli knockout strains lacking trehalose breakdown genes — treB (PTS transporter), treC (trehalose-6-phosphate hydrolase), treA (periplasmic trehalase), and treF (cytoplasmic trehalase) — were obtained from the Keio collection (Baba et al., 2006). Wild-type E. coli BW25113 served as the control strain. Strains were first grown overnight in LB medium at 37 °C with shaking (200 rpm). Cultures were washed twice with M9 minimal medium to remove residual nutrients and subsequently inoculated into fresh M9 minimal medium supplemented with 2% trehalose (w/v), 2 mM amino acids, 2 mM MgSO_4_, and 1 mM CaCl_2_, at an initial optical density of OD_600_ ≈ 0.1. Cultures were incubated at 37 °C with shaking, and OD_600_ was recorded hourly for 24 h using a spectrophotometer. Growth measurements for each strain were normalized to the OD_600_ value at time zero. All growth curves were generated from three independent biological replicates, and the mean ± standard deviation (SD) was plotted.

## Data Availability

All necessary data is available at https://figshare.com/s/04c1bad8d8c6281f1891. Supplementary materials are available to download.

## Acknowledgements

We thank Dr. Anjana Badrinarayanan and Dr. P. Shivaprasad from National Centre for Biological Sciences (NCBS), Bengaluru, India, for helpful discussions, and members of the SL and AS laboratories for insightful feedback throughout the study. This work was supported by the DBT–Wellcome Trust India Alliance Senior Fellowship (IA/S/21/2/505922), the DBT S. Ramachandran National Bioscience Award for Career Development from the Department of Biotechnology, Government of India, and intramural funding to SL; and funding from the Department of Atomic Energy, Government of India (Project Identification No. RTI 4006) to AS.

