## Supplementary Figures for "Evolutionary analysis of Trehalose breakdown pathways"

Supplementary figure 1

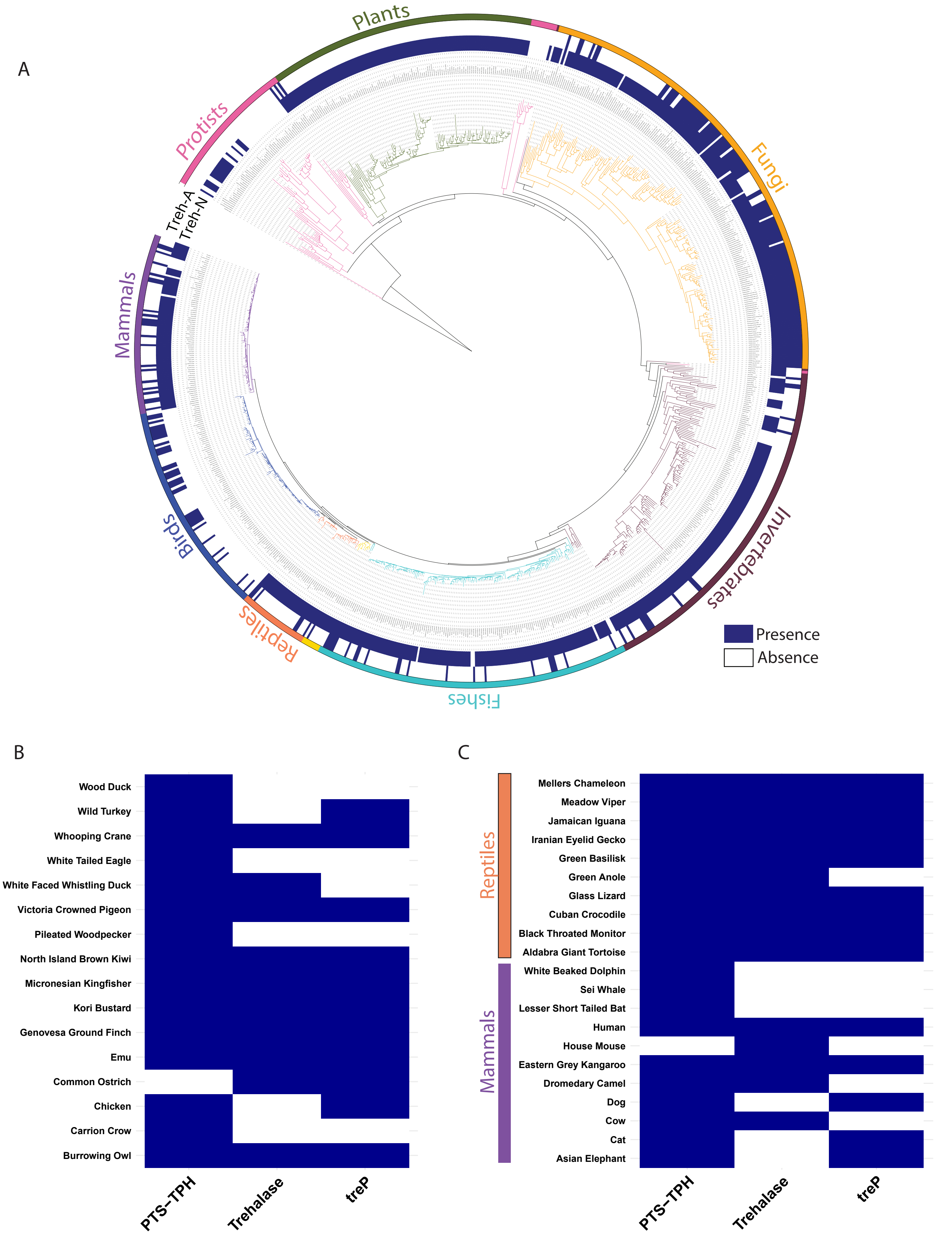

Supplementary figure 2

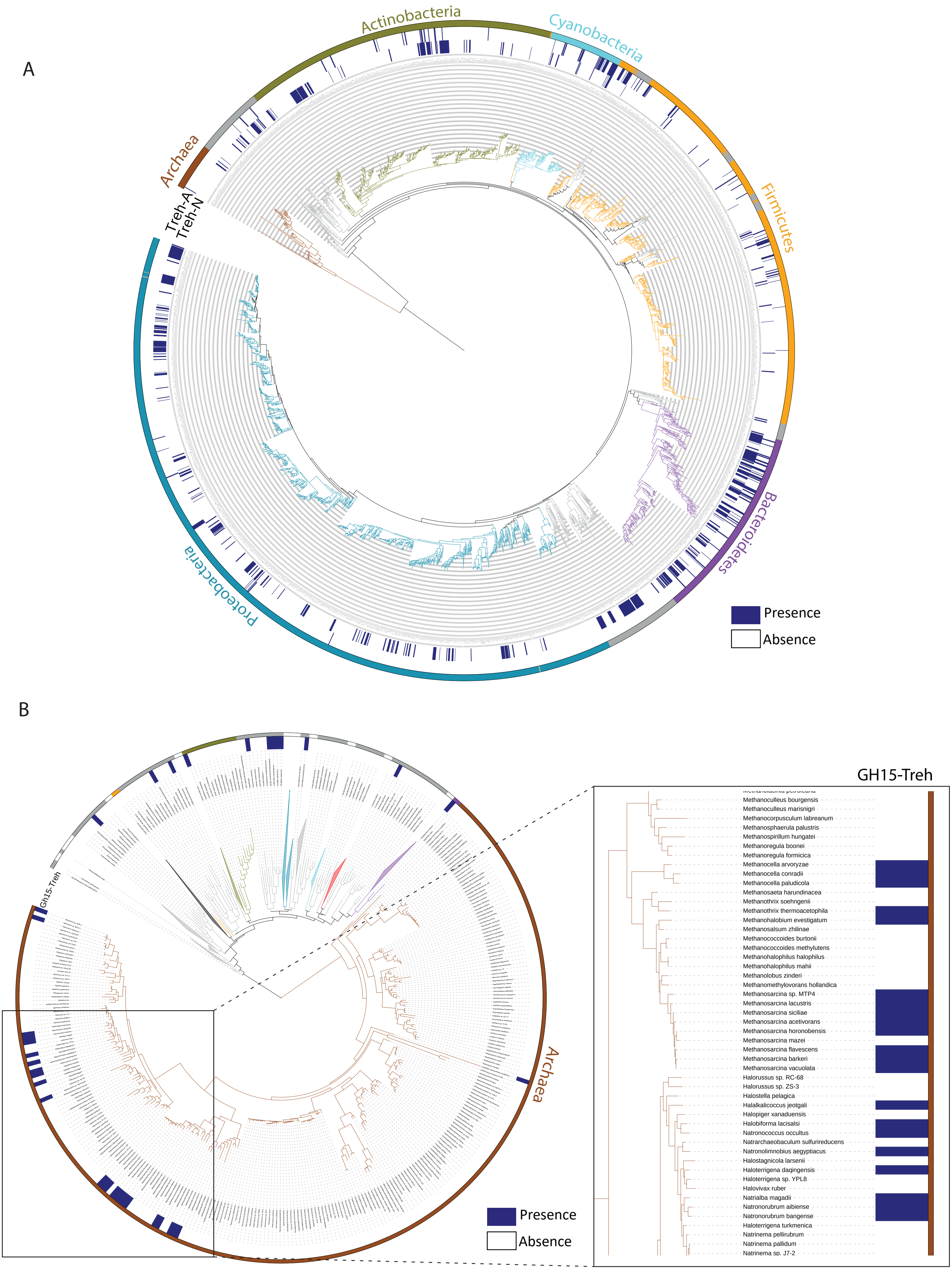

Supplementary figure 3

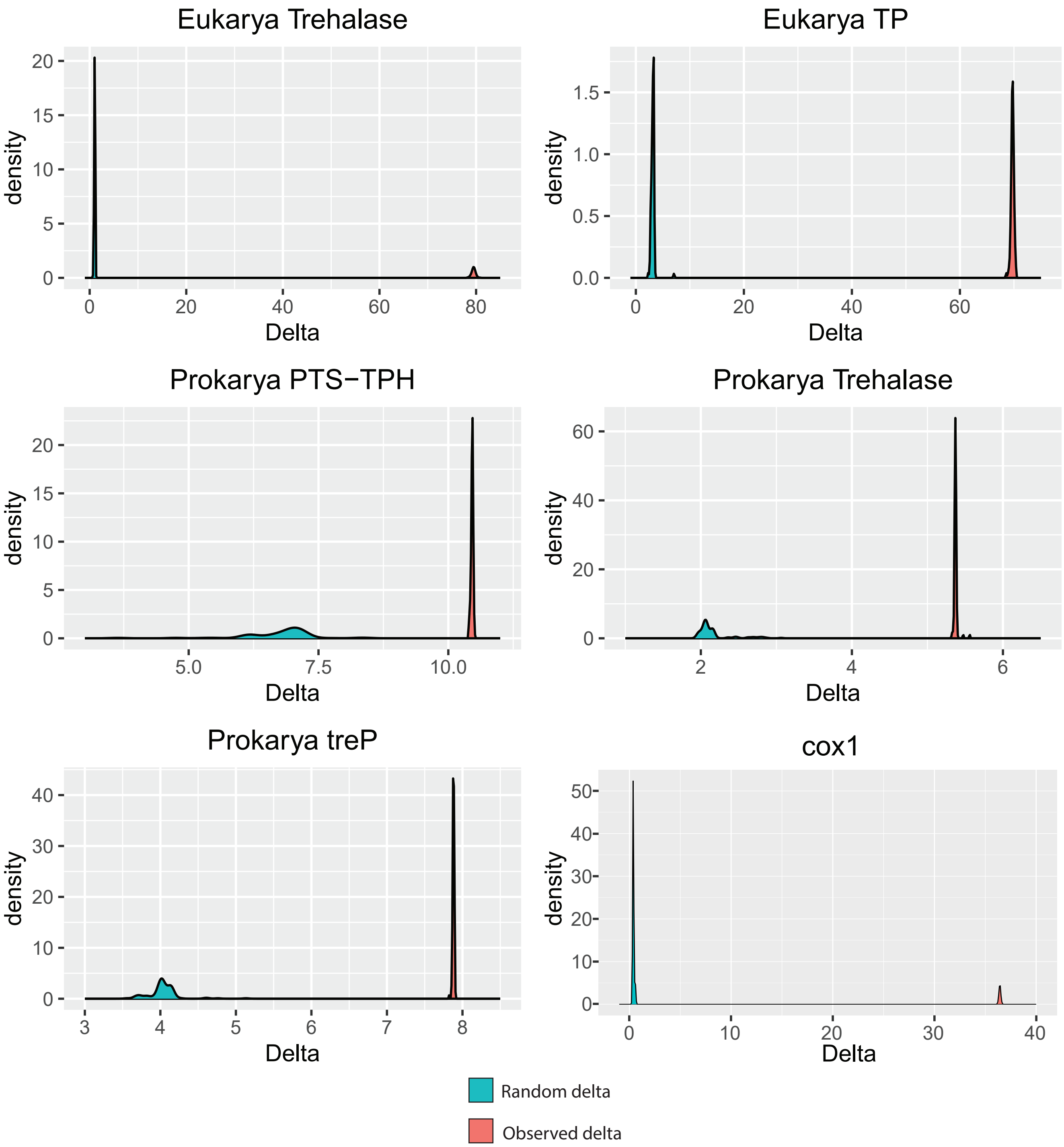

Supplementary figure 4

A

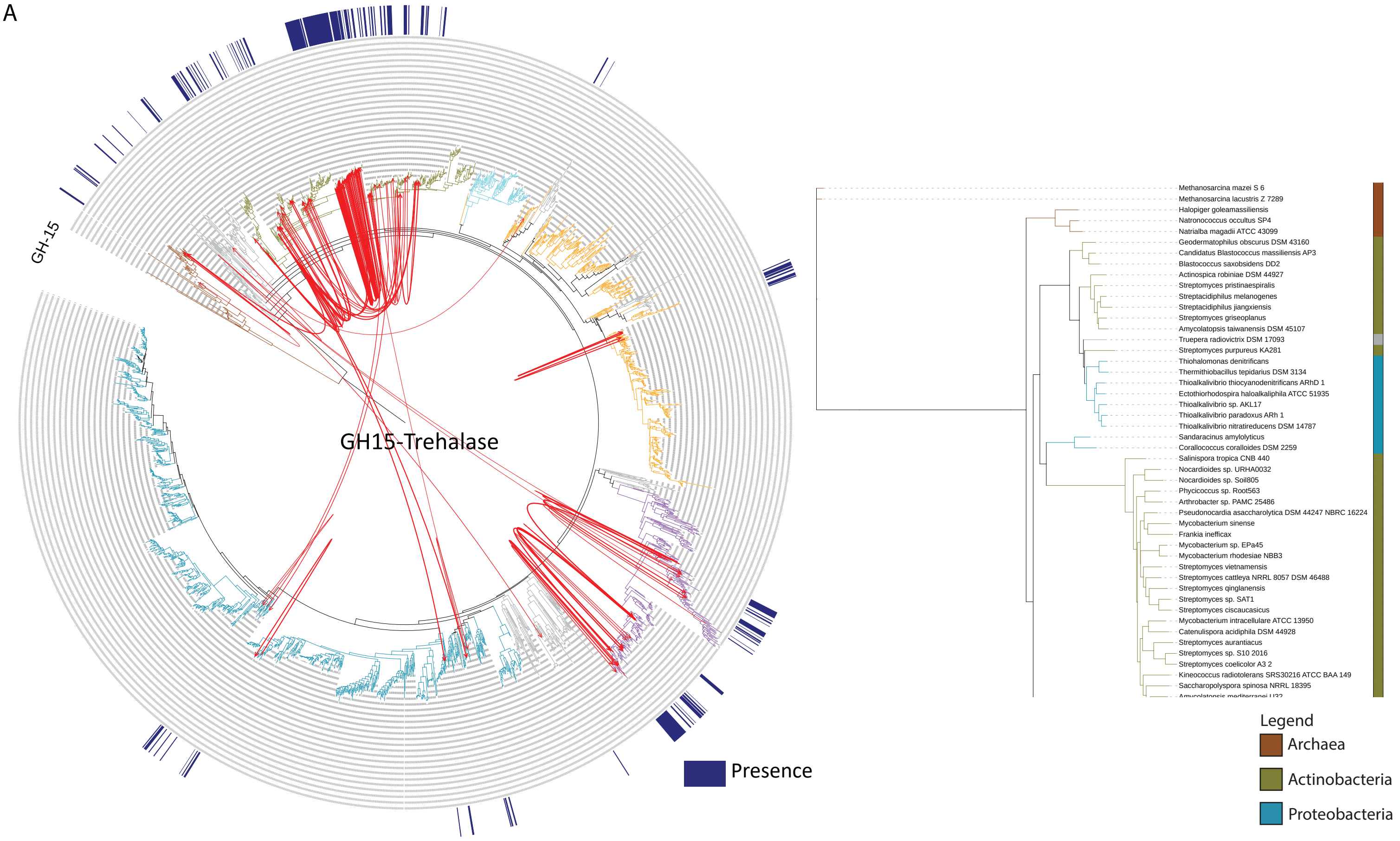


B

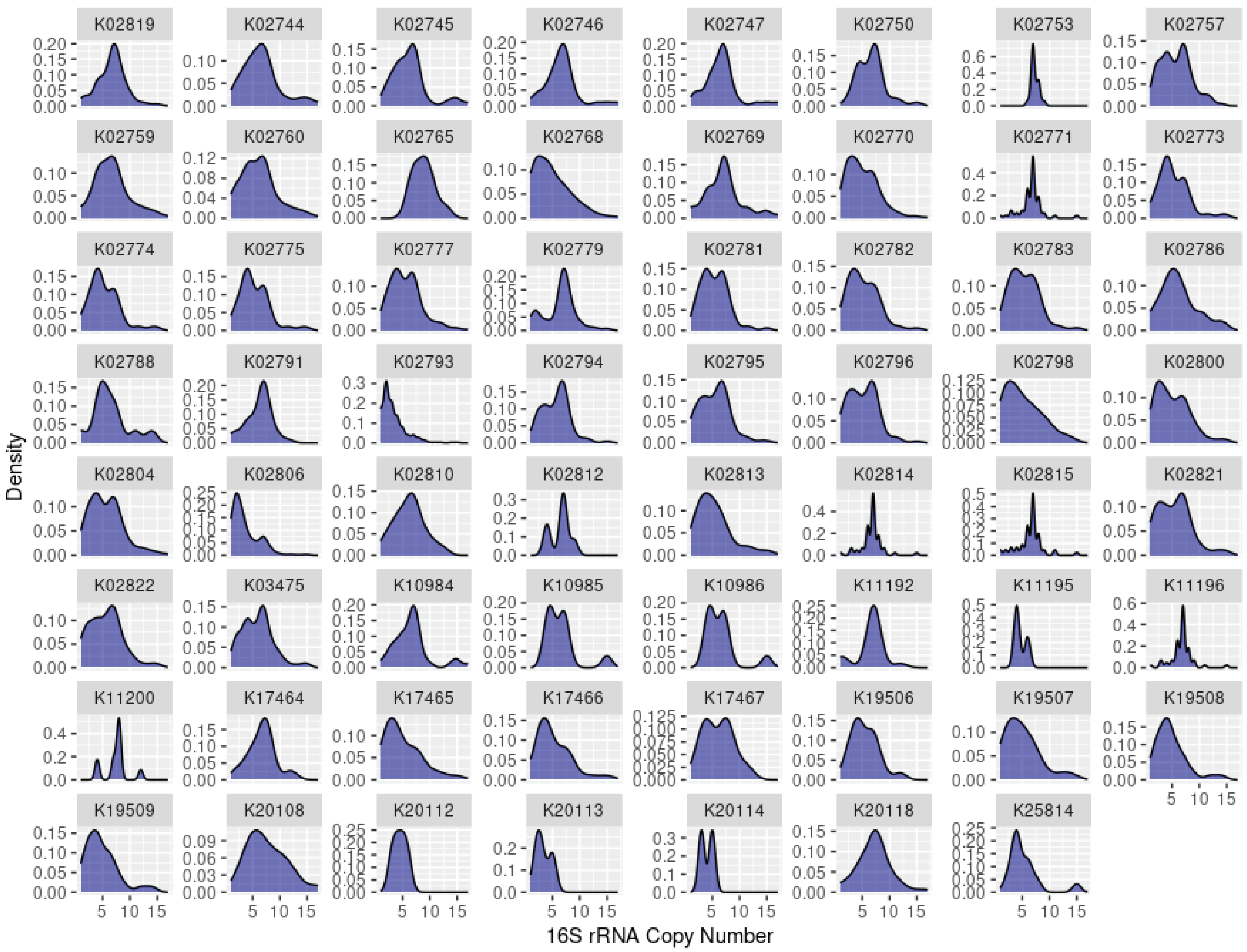

Supplementary figure 5

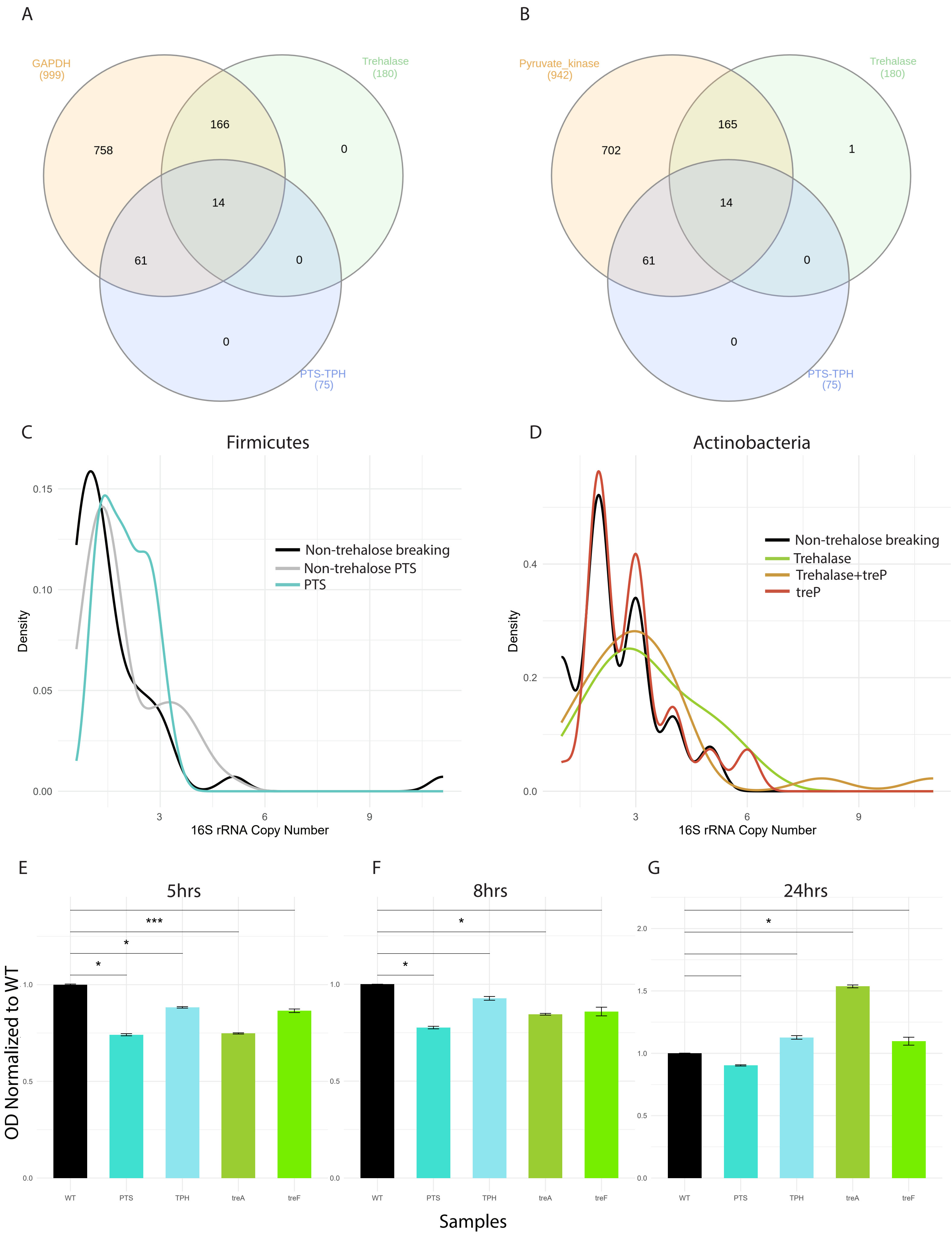

### Supplementary figures legend

**Supplementary Figure 1:**

A. Eukaryotic species phylogeny with a heatmap showing presence (blue) and absence (white) of neutral ($Tre^N$) and acid ($Tre^A$) trehalase across 868 genomes.
B. Heatmap showing the presence of trehalose breakdown pathways (trehalase, PTS-TPH, and treP) in gut microbiota from 15 bird species spanning multiple clades.
C. Comparative heatmap showing the distribution of the same pathways in the gut microbiomes of mammals and reptiles.

**Supplementary Figure 2:**

A. Prokaryotic species phylogeny with a heatmap showing presence (blue) and absence (white) of neutral ($Tre^N$) and acid ($Tre^A$) trehalases.
B. Prokaryotic species phylogeny with a heatmap showing presence (blue) and absence (white) of GH15 trehalase. The bacterial clades are collapsed highlighting its presence in Archaea. Inset figure showing the presence in Methanomicrobia and Halobacteriales.

**Supplementary Figure 3:** Delta statistic values for trehalase, PTS-TPH, and treP pathways compared to the control gene (COX1), indicating weak phylogenetic signal for all trehalose breakdown systems.

**Supplementary Figure 4:**

A. Species tree showing inferred HGT events for GH15 trehalase, with the inset gene tree highlighting cross-phyla clustering consistent with HGT.
B. Distribution of 16S rRNA copy numbers in organisms encoding trehalose-specific PTS versus non-trehalose PTS systems.

**Supplementary Figure 5:**

(A-B) Venn diagrams showing the presence of key glycolytic enzymes—GAPDH (A) and pyruvate kinase (B)—in organisms containing Treh and/or PTS pathways.
(C–D) Distribution of 16S rRNA copy numbers in trehalose-utilizing and non-utilizing Firmicutes (C) and Actinobacteria (D).

(E–G) Relative growth ($OD_{600}$) of *E. coli* knockout strains lacking treB (PTS), treC (trehalose-6-phosphate hydrolase), treA (periplasmic trehalase), or treF (cytoplasmic trehalase), normalized to WT at 5, 8, and 24 hours.
